# Reducing environmentally mediated transmission to moderate impacts of an emerging wildlife disease

**DOI:** 10.1101/2022.07.25.501399

**Authors:** Joseph R. Hoyt, Katy L. Parise, John E. DePue, Heather M. Kaarakka, Jennifer A. Redell, William H. Scullon, Rich O’Reskie, Jeffrey T. Foster, A. Marm Kilpatrick, Kate E. Langwig, J. Paul White

**Author notes:** Corresponding author –.

## Abstract

1. Emerging infectious diseases are a serious threat to wildlife communities, and the ability of pathogens to survive in the environment can exacerbate disease impacts on hosts and increase the likelihood of species extinction. Targeted removal or control of these environmental reservoirs could provide an effective mitigation strategy for reducing disease impacts but is rarely used in wildlife disease control.
2. We examined the effectiveness of managing environmental transmission to reduce impacts of an emerging infectious disease of bats, white-nose syndrome. We used a chemical disinfectant, chlorine dioxide (ClO_2_), to experimentally reduce *Pseudogymnoascus destructans*, the fungal pathogen causing WNS, in the environment. We conducted laboratory experiments followed by three years of field trials at four abandoned mines to determine whether ClO_2_ could effectively reduce *P. destructans* in the environment, reduce host infection, and limit population impacts.
3. ClO_2_ was effective at killing *P. destructans in vitro* across a range of concentrations. In field settings, higher concentrations of ClO_2_ treatment sufficiently reduced viable *P. destructans* conidia in the environment.
4. The reduction in the environmental reservoir at treatment sites resulted in lower fungal loads on bats compared to untreated control populations. Survival following treatment was higher in little brown bats (*Myotis lucifugus*), and trended higher for tricolored bats (*Perimyotis subflavus*) compared to untreated sites.
5. These findings support the management of environmental reservoirs as an effective control strategy for wildlife disease and provide a valuable tool for ongoing conservation efforts. More broadly, these results highlight how the intensity of environmental reservoirs can have cascading impacts on host infection and population declines.

## Introduction

Emerging infectious diseases present a serious threat to biodiversity conservation (van Riper et al. 1986, LaDeau et al. 2007, Skerratt et al. 2007, Langwig et al. 2012, Hollings et al. 2013), and developing successful control measures to reduce population impacts is difficult given the challenges of working with wildlife (McCallum and Dobson 1995, 2002, Hamede et al. 2012, McCallum 2012, Vander Wal et al. 2014, Langwig et al. 2015c). The ability of pathogens to survive outside their hosts can present further complications for successfully managing wildlife diseases (de Castro and Bolker 2005, Rohani et al. 2009, Turner et al. 2016, Hoyt et al. 2020). Free-living pathogen stages can prevent pathogen fade-out, link otherwise disconnected populations (Rohani et al. 2009, Aiello et al. 2016, Hoyt et al. 2018) and prevent recolonization of extirpated sites (de Castro and Bolker 2005). While the presence of environmental reservoirs and indirect transmission are recognized as important attributes of disease systems (King et al. 2008, Breban et al. 2009, Rohani et al. 2009, Hoyt et al. 2020), the contribution and subsequent control of environmental transmission is a largely underutilized control strategy in wildlife disease in comparison to other forms of transmission (i.e. vector, direct contact). If transmission mediated through an environmental reservoir accounts for a substantial fraction of transmission, then environmental decontamination should reduce both infections and host impacts (Colwell et al. 2003, Knutie et al. 2014, Bosch et al. 2015, Martin et al. 2019, Hoyt et al. 2020).

Environmental sterilization and water treatment are hallmarks of disease prevention and control for both agricultural settings and public health (Mintz et al. 1995). For example, environmental surfaces in hospitals are regularly cleaned to reduce the probability of transmission to healthy patients, and water sterilization reduces the likelihood of infection by gastrointestinal parasites like cholera and typhoid (Colwell and Huq 1994, Colwell et al. 2003). Environmental decontamination is also vital to the control of livestock diseases (Gosling 2018, Wales et al. 2021), and air filtration has become important in the control of SARS-CoV-2 transmission (Allen and Ibrahim 2021). However, management of free-living pathogen stages is infrequently targeted for control in wildlife disease despite the potential utility in providing population-scale benefits in already affected areas (Knutie et al. 2014, Martin et al. 2019) or helping prevent pathogen invasion (Bosch et al. 2015). Many recent conservation-relevant wildlife diseases have free-living stages, which include chytridiomycosis (Mitchell et al. 2008, Kilpatrick et al. 2010), chronic wasting disease (Almberg et al. 2011), snake fungal disease (Campbell et al. 2021), avian influenza (Breban et al. 2009), anthrax (Carlson et al. 2018), and white-nose syndrome (WNS) (Lorch et al. 2013b, Hoyt et al. 2020). This highlights the potential for exploring the effectiveness of reducing environmental transmission to promote positive conservation outcomes.

White-nose syndrome, caused by the fungal pathogen, *Pseudogymnoascus destructans*, is responsible for widespread declines in bat populations throughout eastern North America, with several species experiencing >98% declines (Frick et al. 2010, Langwig et al. 2012, Warnecke et al. 2012). WNS has strong seasonal cycles where bats are only affected during winter when they go into hibernation. If bats survive the winter, they can clear infections over summer when they are euthermic (Langwig et al. 2015a). However, *P. destructans* is known to persist for long periods of time on surfaces of hibernacula where bats hibernate (Lorch et al. 2013b, Hoyt et al. 2015), which serve as an environmental reservoir (Lorch et al. 2013a, Lorch et al. 2013b, Hoyt et al. 2020, Hoyt et al. 2021) for the pathogen and the source of reinfection each winter (Langwig et al. 2015a, Hoyt et al. 2018, Hoyt et al. 2020, Hicks et al. 2021).

Following initial pathogen invasion, sites become increasingly contaminated with *P. destructans* and eventually become heavily contaminated (>50% of locations within hibernacula are contaminated; (Hoyt et al. 2020, Hoyt et al. 2021)). When pathogen contamination in the environment is low, exposure is reduced and host mortality is limited (Hoyt et al. 2020). This is likely because *P. destructans* takes a long time to kill its host (∼100 days post-infection; (Lorch et al. 2011, Warnecke et al. 2012)) and bats are able to emerge in the spring and clear infection. Once the environmental reservoir becomes heavily contaminated, bats become more rapidly infected in early winter (between October and mid-November), resulting in a longer period of infection and high mortality before bats can emerge from hibernation. Reducing the extent and abundance of *P. destructans* in the environment could reduce exposure to *P. destructans* and lower mortality from WNS.

At present, there are few tools for managers to reduce the impacts of WNS (Hoyt et al. 2019, Fletcher et al. 2020), and developing control options to reduce the severity of WNS impacts on bats is a top priority (USFWS 2011). Treating hibernacula to reduce *P. destructans* has several advantages as a management tool, including 1) hibernacula can be treated during the summer when bats are absent, which reduces disturbance to bats during winter; and 2) environmental treatment can be done on a larger scale than individual host treatments. Here we present a series of *in vitro* and *in situ* experiments testing the efficacy of environmental treatment at reducing the population level impacts of white-nose syndrome for two highly impacted species of bat, *Myotis lucifugus* and *Perimyotis subflavus*.

## Methods

### Laboratory experiments

We first assessed the effectiveness of ClO_2_ in killing *P. destructans in vitro* using both inoculated culture plates and natural hibernacula substrate (unsterilized rocks from a naturally contaminated mine in Wisconsin). We chose ClO_2_ because it is a strong oxidizing biocide and is a widely used environmental disinfectant in agricultural and public health settings (Rastogi et al. 2010, Al-Otoum et al. 2016). We inoculated Sabouraud dextrose agar (SDA) plates containing Chloramphenicol and Gentamicin (C&G) with 10µl of PBST_20_ solution containing 10^5^ conidia/µl of *P. destructans*. Rock substrate was collected from a mine in Wisconsin and spiked with the same concentration as above. Inoculum was prepared from *P. destructans* cultures that were grown for 30 days at 7°C on SDA with C&G. There were six replicates per concentration for the culture plates and four replicates for the rock substrate. Each replicate was placed in a small environmental chamber (30×15×8 cm plastic containers). We aerosolized ClO_2_ using a mechanical thermal fogger (Vectorfog - C20 1.5L ULV Electric Fogger, South Korea) that produced droplets ∼15 µm in size and had a flow rate up to 0.5 L per minute (30 L per hour). We sprayed three concentrations (100, 250, and 500 ppm) of ClO_2_ or sterile water into the chambers for a total of 10 seconds, and then sealed them and placed them in an incubator at 7°C. ClO_2_ quickly degrades in the air into chlorite and chlorate ions, resulting in contact time for this experiment lasting ∼1 hr. Seven days post-treatment, the *P. destructans-*inoculated rocks were swabbed and plated on SDA. All plates were visually checked every seven days for growth for four weeks. Plates were classified as either growth or no growth following the end of the observation period.

### Field treatment with 500 ppm

Following the laboratory validation that ClO_2_ could kill *P. destructans*, we treated three inactive mines with ClO_2_ in Michigan (MI_BC and MI_TM) and Wisconsin (WI_PS) in 2017 where WNS was detected in the previous year (2015/2016). Sampling bats and the environment at control sites was done as part of a long-term project focused on understanding the invasion and impacts of *P. destructans* on bat communities (Hoyt et al. 2018, Hoyt et al. 2020, Hopkins et al. 2021, Langwig et al. 2021). These data were leveraged as a baseline for comparison against our treated sites to assess differences in pathogen prevalence, fungal loads, and apparent survival. Bats at treatment and control sites were sampled for *P. destructans* twice each winter and were matched by the number of years since *P. destructans* detection because there are strong differences among disease phases (invasion, epidemic, and established) but sites within a phase have similar disease dynamics (Langwig et al. 2015a, Langwig et al. 2015b, Hoyt et al. 2020). Standardized swabs were collected from each bat’s forearm and muzzle, which were stored in RNAlater until samples were tested using qPCR (Muller et al. 2013, Langwig et al. 2015a). Finally, we measured the temperature of each sampled bat using Fluke 62 MAX IR laser thermometer to quantify the temperature of the substrate directly adjacent to the bat (<1 cm).

We treated the environment with ClO_2_ using a licensed professional in late summer, when bats were absent from sites. During the initial treatment, ClO_2_ was applied using a commercial thermal fogger (IGEBA TF-35, Weitnau, Germany) at a concentration of 500 ppm to the entire site with an exposure time of ∼60 min. The ClO_2_ fog allowed for the treatment to reach cracks and crevices where bats may roost but would otherwise be difficult to access by personnel. The entrance of each site was sealed off for 48 h to minimize airflow and restrict access. The barrier on the entrance of the site was then removed and bats were allowed to naturally access sites several months later during fall swarm.

To determine the efficacy of the treatment in the environment, we collected 120 randomly located environmental swab samples before treatment during the summer. Sixty of these locations were marked using reflective tape for a pre- and post-treatment comparison. One to seven days after treatment, 120 locations within the site were sampled, including the 60 previously marked locations. Swabs were tested using qPCR (Muller et al. 2013) to determine the presence and quantity of *P. destructans* pre- and post-treatment. Bats were then sampled in November and March following the treatment to examine differences in fungal prevalence and loads in treated versus control sites.

### Field treatment with 1000ppm

In 2018 and 2019, we performed additional field trials where we modified the treatment and environmental sampling procedure based on results from the 2017 experiment. Treatments were conducted over two years at the same site (WI_GB) where we used a higher concentration (1000 ppm) and contact time (∼2h) based on preliminary results, suggesting that initial treatment with 500 ppm was insufficient to remove *P. destructans* from the environmental locations pre- and post-treatment. In these experiments we also added an assessment for viable *P. destructans* by culturing both pre and post-treatment from the environment. We rubbed a paired swab adjacent to our qPCR sample and streaked this on a plate containing SDA with Chloramphenicol and Gentamicin. Plates were shipped back to the lab on cold packs, placed in the fridge at 5ºC, and allowed to incubate for 30 d to assess *P. destructans* growth. In addition, during this experiment, we applied a unique aluminum forearm band to each bat in early winter at treatment and control sites to determine how many bats that we sampled and marked in November were recaptured in late winter (March). Based on previous research, we assumed that bats that left the hibernaculum before March likely died from WNS, whereas bats that were alive in March survived over the winter (Hoyt et al. 2019, Hopkins et al. 2021). We also calculated survival between March and the following November (between winter survival) to estimate the probability of the bats that survived until March returned to the site the following winter. As before, we visited control and treated sites in early and late winter to collect epidermal swab samples from hibernating bats and the environment to test for the presence and abundance of *P. destructans*.

All work was conducted under relevant federal (#TE64081B_1) and state wildlife scientific permits and with approvals of the IACUC of Virginia Tech (17-180) and the University of California, Santa Cruz (Kilpm1705, Kilpm1509). Wisconsin Department of Agriculture, Trade and Consumer Protections authorized exemption from a WI Experimental Use Permit for GO-2 EPA Reg no 84912-2. The US Environmental Protection Agency authorized an exemption from an Experimental Use Permit. We obtained a WDNR Temporary Access Permit for the Villages of Osceola and Mineral Point and a WDNR Pesticide Pollutant Permit.

### Analyses

Data were analyzed using Bayesian hierarchical models, and we assessed statistical support for effects of treatment using credible intervals that do not overlap zero. We fit all models (unless otherwise noted) using the No-U-Turn Sampler (NUTS), an extension of Hamiltonian Markov chain Monte Carlo (HMCMC). All Bayesian models were created in the Stan computational framework (http://mc-stan.org/) accessed with the “brms” package in program R (Bürkner 2017). To improve convergence and avoid over-fitting, we specified weakly informative priors (a normal distribution with mean of zero and standard deviation of ten), unless otherwise noted. Models were run with a total of 4 chains for 2000 iterations each, with a burn-in period of 1000 iterations per chain resulting in 4000 posterior samples, which, given the more efficient NUTS sampler, was sufficient to achieve adequate mixing and convergence. All 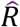 values were less than or equal to 1.01, indicating model convergence.

To estimate the effectiveness of ClO_2_ concentrations at killing *P. destructans* in the laboratory, we used a Bayesian hierarchical model with growth or no growth of *P. destructans* on each plate as our Bernoulli (0/1) response variable, and the concentration of ClO_2_ as our categorical population-level effect (predictors or fixed effects in a frequentist context) with an additive effect for the type of media inoculated (culture plates or rock substrate).

To determine whether *P. destructans* was sufficiently reduced in the environment following field treatment, we initially assessed differences in qPCR fungal loads in the environment pre- and post-treatment with a gaussian distribution and log_10_ fungal loads as our response variable and sampling period (pre- or post-treatment) as our population-level effect. To determine whether there were differences in viable conidia in the environment, we performed an identical analysis as above, but the growth of *P. destructans* on each culture plate was our Bernoulli (0/1) response variable.

To examine *P. destructans* dynamics on bats following treatment, we analyzed the change in *P. destructans* prevalence and fungal loads over winter. For the bat prevalence analyses, our response variable was the detection of *P. destructans* in a sample (0|1), and we included a three-way interaction between sampling date, species (levels = *M. lucifugus* and *P. subflavus*), and treatment (levels = control, 500 ppm ClO_2_, or 1000 ppm ClO_2_) with site as a group level effect. To estimate changes in fungal loads, we ran an identical model structure described above with a gaussian distribution and log_10_ fungal loads as our response variable. Fungal loads are strongly dependent on bat roosting temperature, and warmer sites have faster growth rates (Langwig et al. 2016, Hopkins et al. 2021, Grimaudo et al. 2022) so we selected control sites based on comparable temperature profiles. For the 1000 ppm and 500 ppm treatment comparisons of fungal prevalence and loads overwinter on bats, we included control sites with an average temperature above 7ºC (average control site temperature 9.56±1.52ºC), which is more closely aligned with the treatment site temperature (1000 ppm: WI_GB = 11.0±0.58ºC; 500 ppm: MI_BC = 7.88±1.1ºC, MI_TM = 7.69±0.84 ºC, and WI_PS 10.59±0.26 ºC).

For the field experiment with a 1000 ppm concentration of ClO_2_, we estimated differences in apparent survival between control and treated populations both over and between winters for two species, *M. lucifugus* and *P. subflavus*. Our response variable was whether a bat was recaptured at the end of winter or the following winter for the between winter analysis (0|1), and we included treatment (levels= control or treated) as our population-level effect and site as a group-level effect. WNS impacts on *M. lucifugus* survival also depend strongly on temperature (Johnson et al. 2014, Langwig et al. 2016, Hopkins et al. 2021) so we included temperature as an interaction with treatment for the overwinter analysis. For graphical representation, we extracted the coefficients for the average temperature of our treated sites to show estimated differences at comparable temperatures. Because there was only support for including temperature in the overwinter survival analysis, it was omitted from the *M. lucifugus* between winter analysis and from both overwinter and between winter analyses for *P. subflavus*.

## Results

### Laboratory exposure

Two concentrations of ClO_2_ (250–500 ppm) successfully killed all *P. destructans* colonies on both inoculated culture plates and on natural rock substrate (Figure 1). Growth on plates exposed to 100 ppm of ClO_2_ was reduced in some plates but did not differ from the control plates (results formatted as coefficient ± standard deviation (95% credible intervals); 100 ppm treatment coef: -2.40±2.88 (-8.96, 2.48)), but there was support that *P. destructans* growth was reduced compared to control when exposed to 250 ppm or 500 ppm of ClO_2_ (250 ppm coef: - 14.66±5.22 (-26.42, -6.46), 500 ppm coef: -14.79±5.38 (-27.03, -6.46)). *Pseudogymnoascus destructans* inoculated culture plates and rocks showed similar results, and there was no support for an effect of the rock substrate differing from culture plates across all treatments (Figure 1 right; rock substrate coef: 2.49±2.96 (-2.00, 9.53).

**Figure 1:**
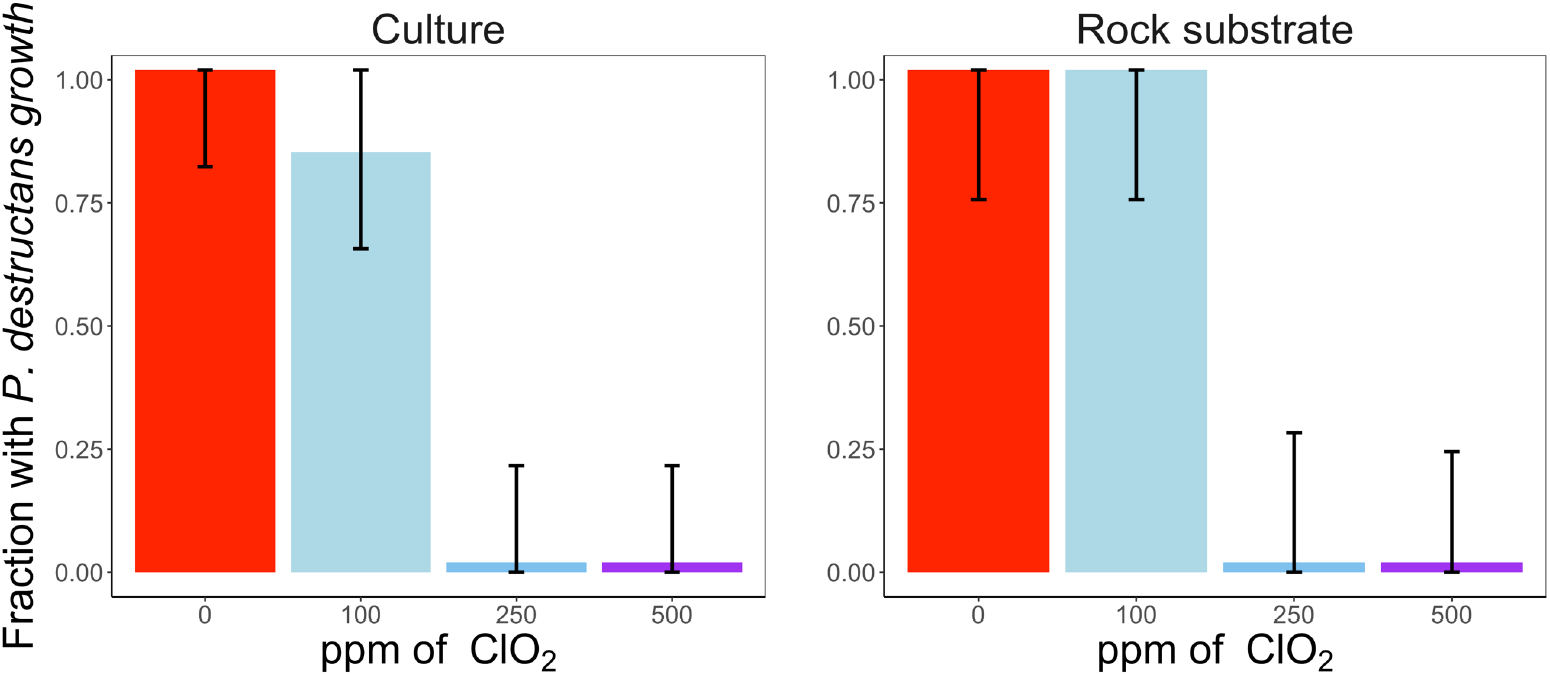
*In vitro P. destructans* challenge experiments with varying concentrations of ClO_2_. Bar plots show the fraction of plates that had *P. destructans* growth following treatment. The error bars show the standard error. Left) shows the fraction of six inoculated plates that had *P. destructans* growth across a range of ClO_2_ concentrations (0, 100, 250 and 500 ppm). Right) shows the fraction of four inoculated rock substrate that had culturable *P. destructans* growth across a range of ClO_2_ concentrations.

### Environmental cleaning effectiveness at reducing the *P. destructans* reservoir

Using qPCR quantification, we found no support for differences in *P. destructans* loads in the environment after treatment with 500 ppm ClO_2_ (Figure 2A; 500 ppm post-treatment coef: 0.04±0.06 (-0.08, 0.17)), or 1000 ppm ClO_2_ (Figure 2B; 1000 ppm post-treatment coef: - 0.02±0.12 (-0.26, 0.22)). In the 1000 ppm ClO_2_ experiments, we tested whether the qPCR was amplifying DNA from non-viable conidia by culturing samples pre and post-treatment to check for viability during the second and third treatment with 1000 ppm ClO_2_ in summers 2018 and 2019. We found statistical support for a reduction in environmental pathogen prevalence following treatment with the 1000 ppm ClO_2_ exposure when measured using culturing of viable fungal conidia (Figure 2C; post-treatment coef: -8.27±4.58 (-19.36, -1.95)). The reduction in prevalence was similar between summers and was combined for this analysis (see Figure S1).

**Figure 2:**
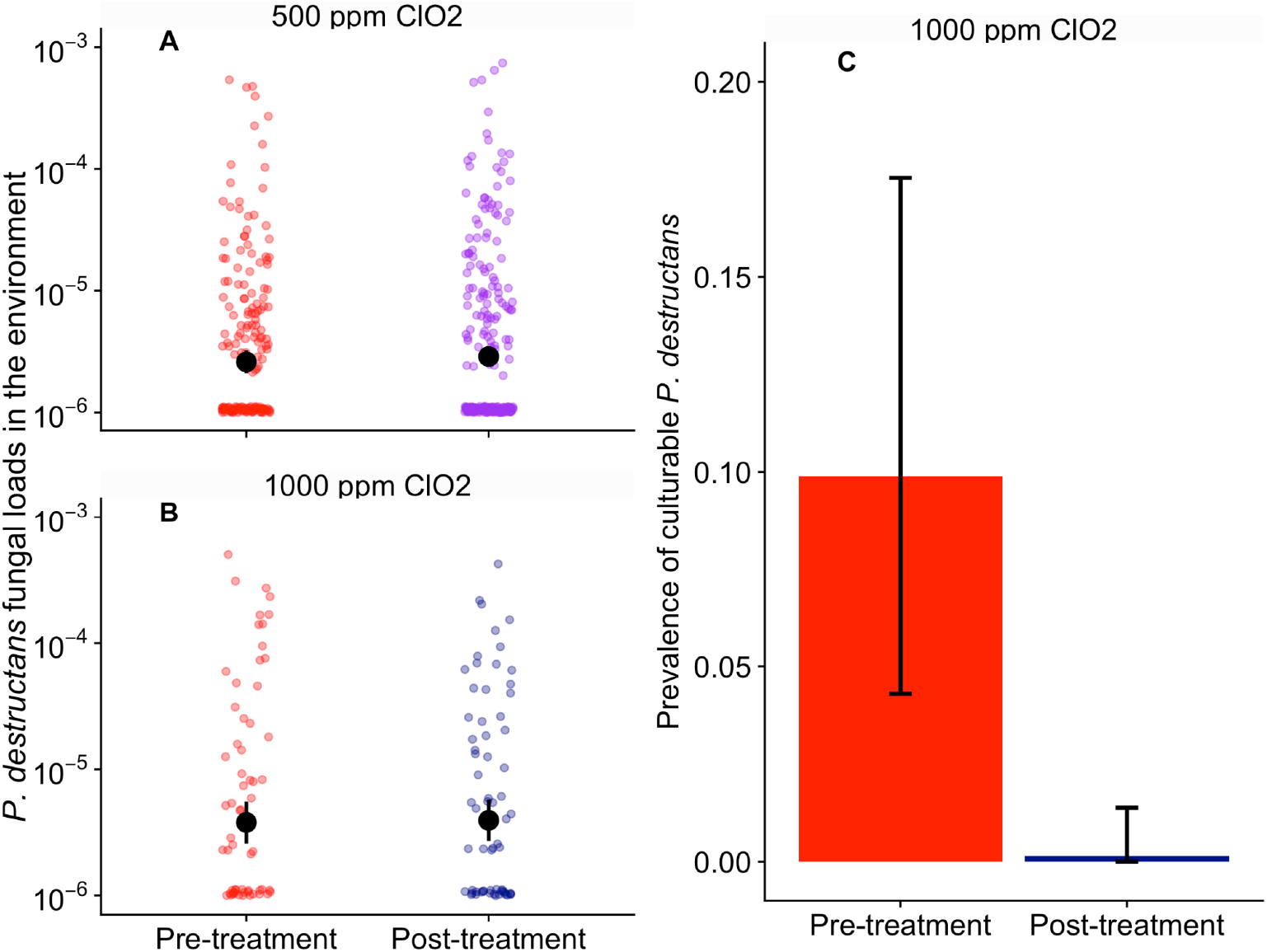
Environmental swab and culture results pre- and post-treatment with ClO_2_ over three years. The colors indicate the pre (red) and post-treatment (purple & blue). Treatment in 2017 was done at three sites (MI_TM, MI_BC, and WI_PS). Treatments in 2018 and 2019 were done at a single site (WI_GB) using a higher concentration (1000 ppm) and a longer exposure period (∼2 h) of ClO_2_. A) Log_10_ fungal loads of *P. destructans* (ng DNA) measured using qPCR pre-treatment (500 ppm: N = 240) and post-treatment (500 ppm: N = 320). Black points show the mean loads and lines show the 95% credible intervals. C) Fraction of swabs with culturable *P. destructans* pre- (1000 ppm: N = 70) and post-treatment (1000 ppm: N = 70). Error bars show the 95% credible intervals and a small constant was added to panel C for visual representation. Cultured samples taken after treatment with 1000 ppm show statistical support for a reduction in viable *P. destructans* following treatment (-8.27±4.58 (-19.36, -1.95)).

### Environmental cleaning effects on host infection

To determine whether environmental decontamination with ClO_2_ successfully reduced host infection intensity, we sampled bats in treated and control sites during early and late winter following treatment. At 1000 ppm, we found statistical support for reductions in pathogen burden at the beginning of hibernation (1.48 and 1.33 orders of magnitude lower for both *M. lucifugus* and *P. subflavus* compared to control sites; Figure 3C; *M. lucifugus* treatment coef: - 4.69±1.45 (-7.43, -1.92); Figure 3D; *P. subflavus* treatment coef: -6.06±1.75 (-9.5, -2.59)). We did not detect an effect of treatment on pathogen prevalence on bats during early winter (Figure 3A; *M. lucifugus* treatment coef: -0.44 ±3.28 (-7.09, 5.68); Fig. 3B; *P. subflavus* treatment coef: - 0.52±3.21 (-7.23, 5.51)). *Myotis lucifugus* at sites that were treated with 500 ppm ClO_2_ showed no reduction in fungal prevalence or fungal loads at the beginning of winter compared with bats at untreated control sites (Figure S2A & B; Table S3 and S4, respectively). Other bat species were not present in sufficient numbers to analyze the effect of treatment on infection.

**Figure 3:**
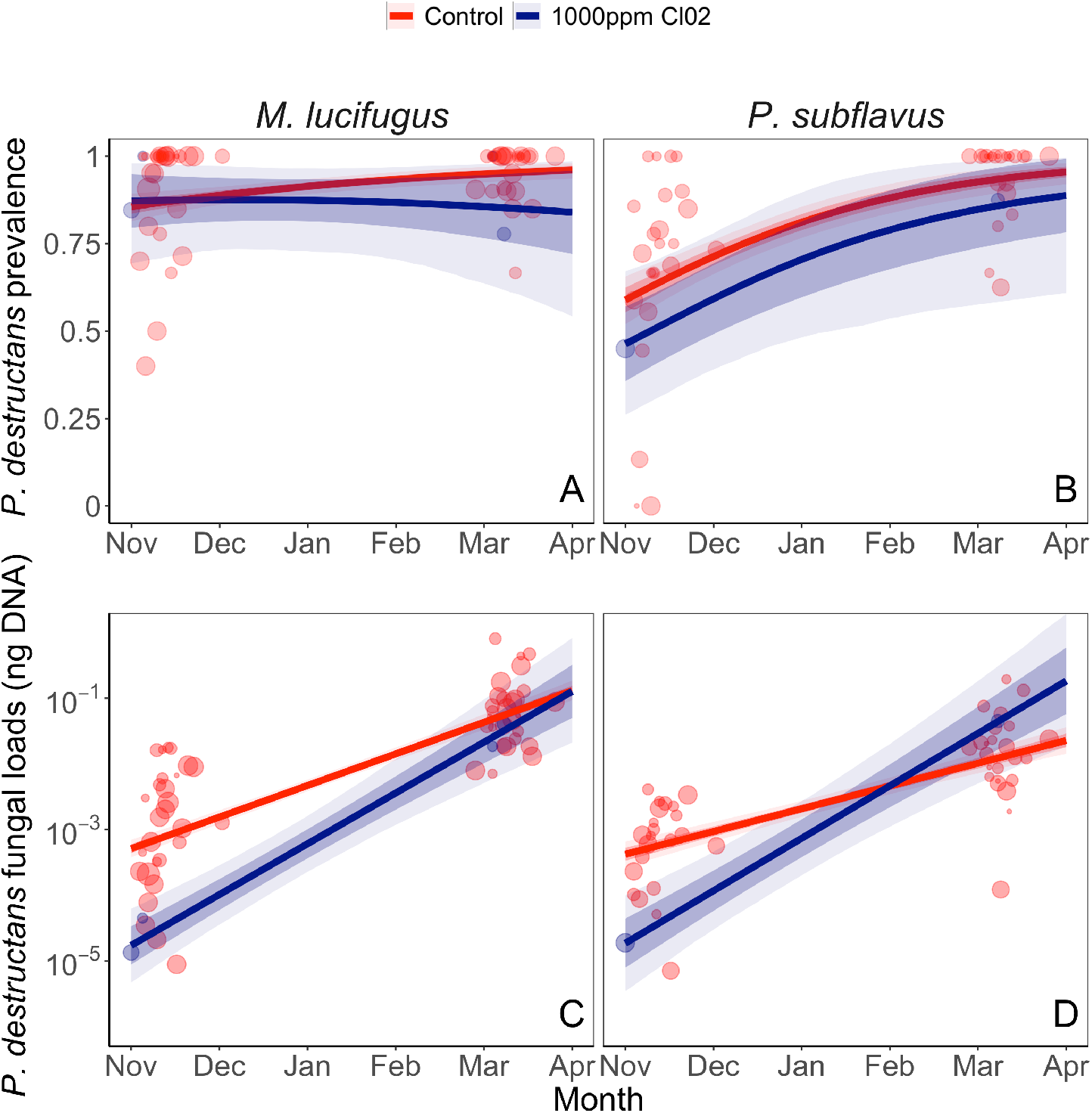
Change in the prevalence and quantity of *P. destructans* DNA on bats in control (red) and ClO_2_ treated (blue) sites. The colored points in each panel show the prevalence for a species at a given sampling event (A & B) or individual fungal loads in early/late winter (C & D) the light-shaded ribbons show the 95% confidence intervals and the darker shaded ribbons indicate the standard deviation. The smaller semi-transparent points show average fungal loads observed for each site. Environmental treatment with 1000ppm ClO_2_ showed statistical support for a reduction in fungal loads for two species, *M. lucifugus* (treatment coef: -4.69±1.45 (-7.43, - 1.92)) and *P. subflavus* (treatment coef: -6.06±1.75 (-9.5, -2.59)).

### Environmental cleaning effects on host survival

To determine whether environmental cleaning increased host survival we compared the probability of recapturing bats in treated versus control sites over winter and between winters. We found statistical support for a higher probability of recapturing *M. lucifugus* following ClO_2_ environmental treatment (1000 ppm, Figure 4A-B). Treatment with 1000 ppm ClO_2_ increased overwinter survival of *M. lucifugus* by 50% from 19% to 69% (Figure 4A; treatment survival coef: 2.37±0.64 (1.14, 3.67)) and between-winter survival by 26% from 20% to 46% (Figure 4B; treatment survival coef: 1.17±0.58 (0.02, 2.28)) compared to control sites. Survival of *P. subflavus* also trended higher, but the 10% (35% in treated and 25% in control) and 12% (28% in treated and 16% in control) differences in survival between treated and control sites within and between winters, respectively, had 95% credible intervals that overlapped zero (Figure 4C; overwinter treatment survival coef: 0.50±0.62 (-0.79, 1.69);Fig 4D; between winter treatment survival coef: 0.67±0.50 (-0.37, 1.60)).

**Figure 4:**
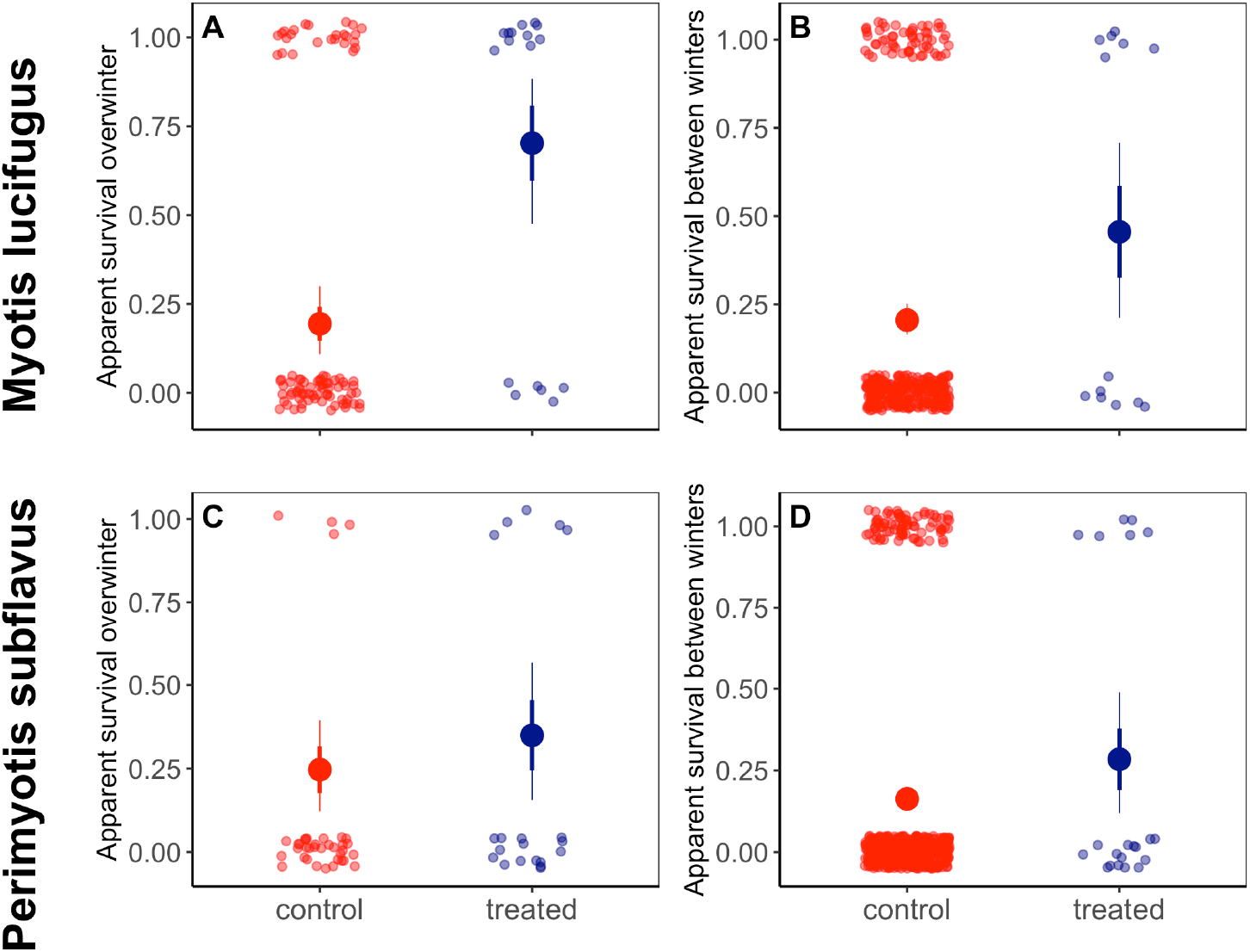
Probability of recapturing *M. lucifugus* or *P. subflavus* at treatment and control sites. Color indicates treatment (blue) and control (red) sites. Large solid points show the model fit, thick lines represent the standard deviation, and thin lines represent the 95% CI. Apparent survival for *M. lucifugus* overwinter (A) and between winters (B) in sites where the environmental reservoir had been reduced using ClO_2_ treatment compared to control sites. Reducing the environmental burden increased the probability of recapturing *M. lucifugus* overwinter (A) by 51% (treatment coef: 1.99±0.33 (1.39, 2.66)) and between winters (B) by 25% (treatment coef: 1.18±0.58 (0.00, 2.66)). There was no statistical support for differences in survival for *P. subflavus* in sites decontaminated with ClO_2_ either overwinter (C) or between winters (D)

## Discussion

We found that reducing the environmental reservoir within hibernacula limited exposure of bats to *P. destructans* and reduced WNS impacts. Higher concentrations of ClO_2_ appeared to kill sufficient amounts of the pathogen in the environment to reduce host infection and increase host survival. More broadly, this study highlights the potential for environmental reservoir control as a tool for wildlife disease management and the links between environmental reservoirs and population-scale impacts.

Disentangling transmission between hosts versus transmission from an environmental reservoir can be exceedingly challenging. Previous research has suggested that the timing of transmission is critical in determining the extent of host impacts for WNS (Hoyt et al. 2020). Sites with environmental pathogen prevalence below 20% in early winter showed stable or growing population growth rates, on average, while sites with higher levels of contamination in the environment had declining populations. Our results provide experimental support for the link between the environmental reservoir and population impacts. Through an experimental reduction in the environmental reservoir we observed reduced infections (as measured by fungal loads), and an increase in apparent survival of bats.

During both treatments, we saw no differences in fungal loads in the environment measured through qPCR before and after environmental decontamination (Figure 2A-B). This is likely because ClO_2_ treatment at 500 ppm or at 1000 ppm did not sufficiently degrade *P. destructans* DNA over the short sampling time (sampling at 3 h, 24 h, or 7 d post-treatment) between treatment and post-treatment sampling and the *P. destructans* specific qPCR still amplified small fragments of DNA from dead fungal cells. Culturing *P. destructans* from the environment did reveal an effect of treatment on *P. destructans* environmental prevalence (Figure 2C), which indicates that treatments were effective at reducing *P. destructans* in the environment. While qPCR is a valuable tool for measuring changes in host infection and environmental reservoir dynamics (Langwig et al. 2015a, Langwig et al. 2016, Hoyt et al. 2020, Hopkins et al. 2021), the short sampling time between treatment and post-treatment sampling suggests that it may not be sufficient for fungal DNA from cells to degrade, and limited our ability to use qPCR to determine whether environmental treatment reduced fungal contamination. Future studies should consider culturing to determine treatment effectiveness or other methods for distinguishing viable and non-viable spores in a sample (Grela et al. 2018). We saw fairly low culture success (∼10% of samples were positive pre-treatment), which may be a result of random marked location sampling where very low amounts of fungus were present and exacerbated difficulties of transferring the pathogen from the hibernacula wall to a swab and then onto a culture plate. Culturing efficacy could be improved by targeting environmental locations that bats use regularly and should have higher concentrations of pathogen.

While we did not perform culturing before and after treatment during the 500 ppm ClO_2_ treatment, the high fungal loads on bats (Figure 3A & C) suggests that this treatment did not kill enough *P. destructans* in the environment to reduce infection in bats. In subsequent experiments using 1000 ppm ClO_2_, where viable *P. destructans* in the environment was clearly reduced, fungal loads on bats were lower in early winter and bats had higher survival, which further supports that the initial treatment concentration was insufficient to reduce fungal burdens in the environment.

Although *P. destructans* loads were lower on bats in early winter following the 1000ppm treatment than at control sites (Figure 3), prevalence and fungal loads eventually reached levels equivalent to control sites and in some cases, even a little higher by the end of winter (Figure 3c-d). This is likely a survival bias due to mortality at the control sites where bats with higher loads in early winter are more likely to die and aren’t sampled during March, which has been shown in previous studies (Langwig et al. 2016, Hopkins et al. 2021). This sampling bias leads to a left-skewed load distribution and reduces the measured late winter fungal loads in bats at control sites (Langwig et al. 2016, Hopkins et al. 2021). We also found that prevalence did not differ between control and treated sites for any of the treatments. This may be due to environmental hotspots of *P. destructans* that bats use within these sites (Hoyt et al. 2018). Even small amounts of viable conidia may be enough to initiate infection, and given the *P. destructans* bat prevalence in early winter typically exceeds 85%, we did not expect to reduce pathogen prevalence on bats. However, the goal of managing environmental reservoirs is not to completely eliminate the pathogen from sites but to simply reduce environmental exposure to *P. destructans*, reducing pathogen loads and allowing bats to survive through winter hibernation.

We observed higher overwinter survival in *M. lucifugus*, but the fraction of marked bats that returned the following winter was ∼50% of the previous year’s late winter count following treatment. This indicates that while environmental decontamination increased within winter survival, a substantial fraction of bats likely did not survive during spring emergence. Following spring emergence, bats infected with *P. destructans* can suffer considerable immune-mediated damage and in some cases suffer a form of immune reconstitution inflammatory syndrome (Meteyer et al. 2012) as their immune systems are reactivated and attack fungal cells that have invaded their wing tissue (Fuller et al. 2020). We still saw a positive effect on survival between winters in our treated site compared to controls but given the loss of ∼50% of individuals that survived over winter, future studies examining the effectiveness of treatments should consider not only differences in survival of bats within a winter but the fraction of bats surviving to the following winter.

There have been concerns over the possible proliferation of *P. destructans* in the absence of competitors or unintended consequences of environmental cleaning on other subterranean organisms. We found no evidence that *P. destructans* amplified in the environment following our treatment, as environmental contamination levels measured through qPCR were similar in control and treated sites. Our sites were chosen because they are inactive mining adits or cellars dug for lagering beer and have been abandoned for <100 years and contain no sensitive species. Cleaning the environment via broad-based disinfectants is best suited for these types of artificial hibernacula that provide little ecological value to species other than bats. Fortunately, many of the largest bat populations in WNS affected areas are found in artificial mines, tunnels, and culverts (Wilder et al. 2011, Kurta and Smith 2014).

In this study, we tested the effects of ClO_2_ treatment on the environmental reservoir of *P. destructans*. ClO_2_ is a powerful oxidizing biocide, is widely used for other fungal pathogens including Anthrax, and can easily be converted to vapor allowing it to reach challenging spaces like cracks and crevices in mines (Rastogi et al. 2010). While this chemical has advantages, there are likely other cleaning methods that may be effective at reducing *P. destructans* in the environment. Alternatives include active (power washing or targeted spraying) or passive removal through chemical and chemical-free inactivation (PEG, VOCs, or UV sterilization (Cornelison et al. 2014, Palmer et al. 2018)). Alternative treatments should consider effectiveness in reaching large portions of the site, effectiveness underground, and scalability. We found that ClO_2_ was effective *in vitro* at low concentrations (250 ppm). However, field application required a much higher concentration to achieve similar outcomes possibly due to numerous unforeseen factors, including site temperature, substrate chemistry, and changing airflow, which likely affected chemical exposure time. Mechanical removal of *P. destructans* could avoid some of these issues, but will be difficult or impossible to implement in larger sites.

Controlling wildlife disease is a critical challenge for resource managers. Conventional methods such as topical host treatments, vaccines, and probiotics are often pursued first for reducing population impacts of disease. This study highlights how understanding the ecological principles underlying disease dynamics can facilitate innovative approaches to managing disease outcomes. Reduced environmental transmission is thought to be one of the factors contributing to lower impacts ofWNS in bats in Europe and Asia, where a natural reduction in the environmental reservoir occurs each summer (Hoyt et al. 2020). Artificial replication of this process (reducing the environmental reservoir) in a region experiencing epizootic declines increased host survival. Finally, understanding environmentally-mediated transmission and its contribution to disease outbreaks is critical for reducing epidemic potential. Future characterization of the environmental reservoir across space and time will provide more accurate and targeted control efforts to break chains of transmission and prevent disease outbreaks.

## Author contributions

JRH and JPW conceived the ideas, JRH, KEL, JED, RO, JTF, AMK and JPW designed methodology; JRH, HMK, JAR, JED, WHS, RO, AMK, KEL, and JPW collected the data; JRH and KLP conducted laboratory procedures; JRH and KEL analyzed the data; JRH led the writing of the manuscript. All authors contributed critically to the drafts and gave final approval for publication.

## Acknowledgements

We would like to acknowledge the generous landowners for providing access to sites for sampling, and the villages of Osceola and Mineral point for their support of the project. We also acknowledge David Blehert, Jeffery Lorch, Michelle Verant, Donald Sockett, and Melissa Behr for their previous contributions to this work. Funding was provided by the joint NSF-NIH-NIFA Ecology and Evolution of Infectious Disease award DEB-1911853, Bat Conservation International and The Nature Conservancy.

## Data availability statement

Data will be made available through the Dryad Digital Repository before publication or upon reviewer request.

