## Supplemental information for "Reducing environmentally mediated transmission to moderate impacts of an emerging wildlife disease"

### Figures and Tables:

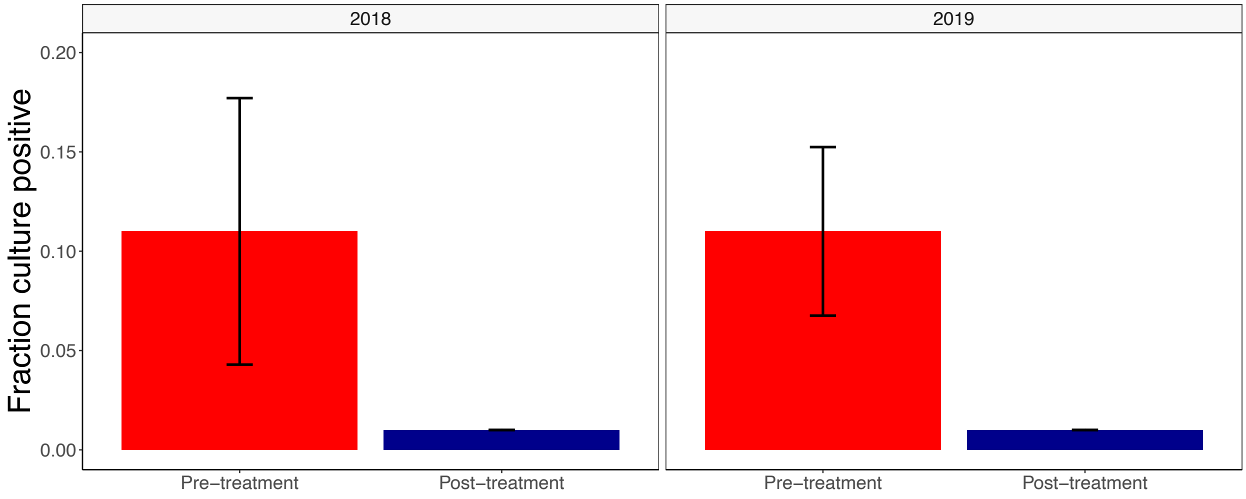


**Figure S1: Culturing results before and after treatment for two treatments at 1000ppm $ClO_2$.** Bars show the number of plates that were culture positive for *P. destructans* before and after treatment. A small constant was added for visualization purposes. Results are combined in Figure 2.

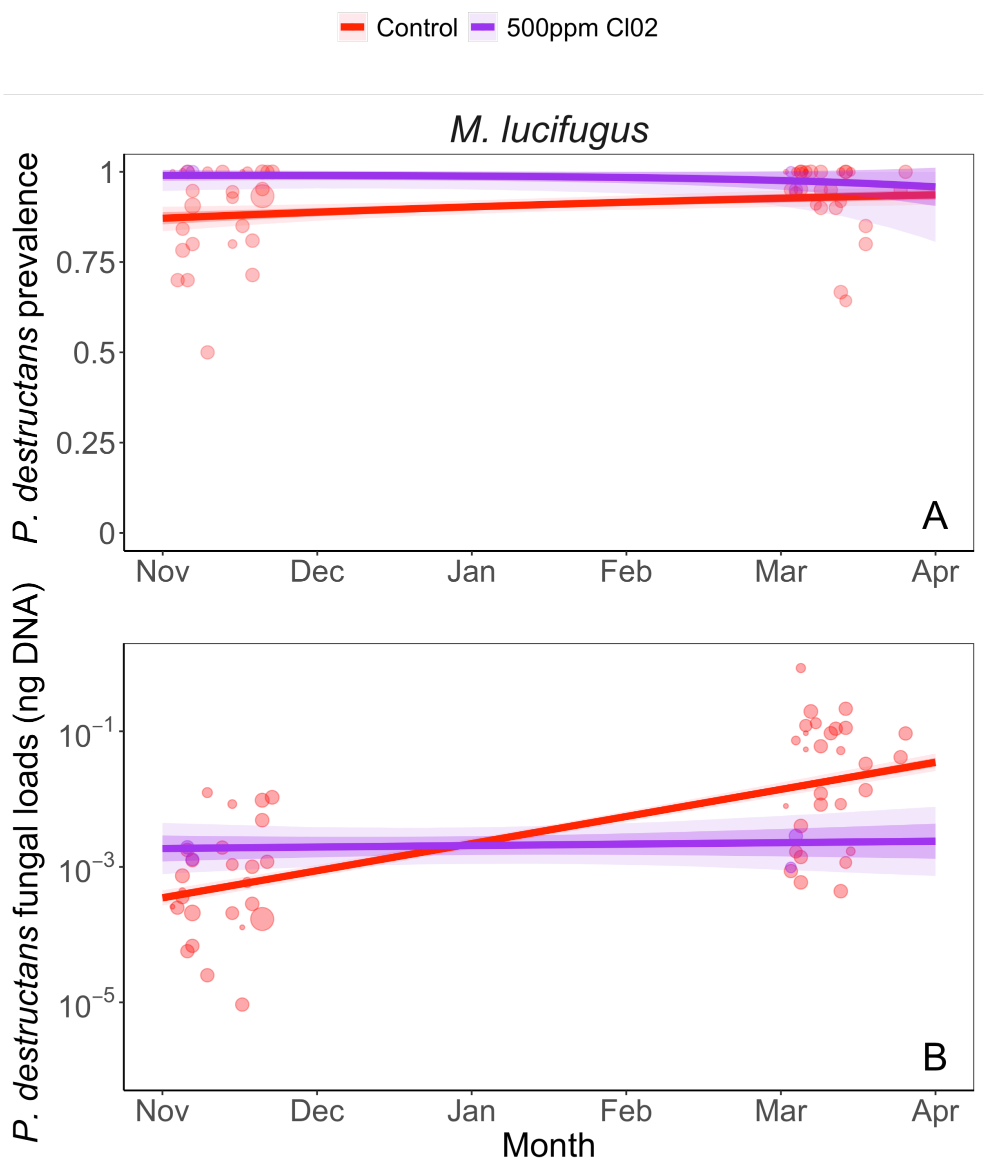


**Figure S2: Fungal prevalence and loads for *M. lucifugus* in sites treated with 500ppm $ClO_2$.** The points in each panel show the prevalence for a species at a given sampling event (A) or avergae fungal loads in early/late winter (B). The light-shaded ribbons show the 95% confidence intervals and the darker shaded ribbons indicate the standard deviation. The smaller semi-transparent points show average fungal loads observed for each site. Statistical tables are contained in Table S3 & S4.

**Table S1: Model output for comparison of bat prevalence in control and treated sites.** Model output from Bayesian hierarchical model with fungal prevalence as our Bernoulli response variable (0|1), a triple interaction between date, species and treatment. Models we run with 4 chains, each with 2000 iterations and a warmup of 1000. Coefficients are shown in relation to the reference (*M. lucifugus* (MYLU) in control sites).

| **Population-level effects** | **Estimate** | **SD** | **L-95% CI** | **U-95% CI** |
|---|---|---|---|---|
| Intercept (MYLU: Control) | -5.71 | 1.03 | -7.85 | -3.73 |
| Date | 0.55 | 0.09 | 0.39 | 0.73 |
| Species-PESU | 4.3 | 1.37 | 1.57 | 7.01 |
| Treatment (1000ppm) | -0.44 | 3.28 | -7.09 | 5.68 |
| Date: species-PESU | -0.26 | 0.11 | -0.48 | -0.04 |
| Date: Treatment (1000ppm) | -0.01 | 0.28 | -0.53 | 0.56 |
| Species-PESU: Treatment (1000ppm) | 4.32 | 4.62 | -4.53 | 13.63 |
| Date: species-PESU: Treatment (1000ppm) | -0.32 | 0.37 | -1.07 | 0.41 |

**Table S2: Model output for comparison of fungal loads on bats in control and treated sites.** Model output from Bayesian hierarchical model with fungal loads as our response, a triple interaction between date, species and treatment. Models we run with 4 chains, each with 2000 iterations and a warmup of 1000. Coefficients are shown in relation to the reference (*M. lucifugus* (MYLU) in control sites).

| **Population-level effects** | **Estimate** | **SD** | **L-95% CI** | **U-95% CI** |
|---|---|---|---|---|
| Intercept (MYLU: Control) | -8.57 | 0.3 | -9.16 | -7.97 |
| Date | 0.48 | 0.02 | 0.44 | 0.52 |
| Species-PESU | 1.36 | 0.53 | 0.34 | 2.42 |
| Treatment (1000ppm) | -4.69 | 1.45 | -7.43 | -1.92 |
| Date: species-PESU | -0.13 | 0.04 | -0.21 | -0.06 |
| Date: Treatment (1000ppm) | 0.29 | 0.11 | 0.07 | 0.5 |
| Species-PESU: Treatment (1000ppm) | -1.62 | 2.32 | -6.26 | 2.94 |
| Date: species-PESU: Treatment (1000ppm) | 0.16 | 0.18 | -0.19 | 0.52 |

**Table S3: Model output for comparison of pathogen prevalence for *M. lucifugus* in control and treated sites (500 ppm).** Model output from Bayesian hierarchical model with fungal prevalence as our bernouli response variable (0|1), a two-way interaction between date and treatment. Models we run with 4 chains, each with 2000 iterations and a warmup of 1000. Coefficients are shown in relation to the reference (control sites).

| **Population-level effects** | **Estimate** | **SD** | **L-95% CI** | **U-95% CI** |
|---|---|---|---|---|
| Intercept (Control) | -1.38 | 0.95 | -3.23 | 0.48 |
| Date | 0.29 | 0.08 | 0.14 | 0.44 |
| Treatment (500ppm) | 9.27 | 6.76 | -2.89 | 23.71 |
| Date: Treatment (500ppm) | -0.55 | 0.49 | -1.55 | 0.39 |

**Table S4: Model output for comparison of fungal loads on *M. lucifugus* in control and treated sites (500 ppm).** Model output from Bayesian hierarchical model with fungal loads as our response, a two-way interaction between date and treatment. Models we run with 4 chains, each with 2000 iterations and a warmup of 1000. Coefficients are shown in relation to the reference (control sites).

| **Population-level effects** | **Estimate** | **SD** | **L-95% CI** | **U-95% CI** |
|---|---|---|---|---|
| Intercept (Control) | -8.59 | 0.29 | -9.14 | -8.01 |
| Date | 0.48 | 0.02 | 0.44 | 0.52 |
| Treatment (500ppm) | 5.16 | 1.18 | 2.8 | 7.52 |
| Date: Treatment (500ppm) | -0.42 | 0.09 | -0.6 | -0.24 |
